# Hematopoietic stem cells fail to regenerate following inflammatory challenge

**DOI:** 10.1101/2020.08.01.230433

**Authors:** Ruzhica Bogeska, Paul Kaschutnig, Malak Fawaz, Ana-Matea Mikecin, Marleen Büchler-Schäff, Stella Paffenholz, Noboru Asada, Felix Frauhammer, Florian Buettner, Melanie Ball, Julia Knoch, Sina Stäble, Dagmar Walter, Amelie Petri, Martha J. Carreño-Gonzalez, Vinona Wagner, Benedikt Brors, Simon Haas, Daniel B. Lipka, Marieke A.G. Essers, Tim Holland-Letz, Jan-Philipp Mallm, Karsten Rippe, Paul S. Frenette, Michael A. Rieger, Michael D. Milsom

## Abstract

Hematopoietic stem cells (HSCs) are canonically defined by their capacity to maintain the HSC pool via self-renewal divisions. However, accumulating evidence suggests that HSC function is instead preserved by sustaining long-term quiescence. Here, we study the kinetics of HSC recovery in mice, following an inflammatory challenge that induces HSCs to exit dormancy. Repeated inflammatory challenge resulted in a progressive depletion of functional HSCs, with no sign of later recovery. Underlying this observation, label retention experiments demonstrated that self-renewal divisions were absent or extremely rare during challenge, as well as during any subsequent recovery period. While depletion of functional HSCs held no immediate consequences, young mice exposed to inflammatory challenge developed blood and bone marrow hypocellularity in old age, similar to elderly humans. The progressive, irreversible attrition of HSC function demonstrates that discreet instances of inflammatory stress can have an irreversible and therefore cumulative impact on HSC function, even when separated by several months. These findings have important implications for our understanding of the role of inflammation as a mediator of dysfunctional tissue maintenance and regeneration during ageing.

## Main

Within regenerating tissues, adult stem cells are thought to comprise a life-long reserve from which mature cells are ultimately replenished in response to normal turnover or following depletion resulting from injury. The critical concept underlying this hypothesis is that adult stem cells are thought to possess extensive self-renewal potential *in vivo*, being capable of reversibly switching from quiescence to active proliferation in order to sustain hematopoiesis (Wilson et al., 2008). In such a setting, the functional status of stem cells would be preserved, regardless of the degree to which they had contributed to production of mature cells. However, such a scenario is not compatible with the popular notion that the progressive attrition of stem cell function is an important root cause of a number of diseases, particularly in the context of ageing (Lopez-Otin et al., 2013).

In the mammalian blood system, HSCs typically exist in a quiescent state and are thought to infrequently contribute towards mature blood cell production under homeostatic conditions (Busch et al., 2015; Cheshier et al., 1999; Sun et al., 2014). Label retention experiments in the laboratory mouse illustrate the heterogeneous proliferative behavior of native HSCs and demonstrate that so-called “dormant” HSCs, which possess a comparatively low division history, retain the highest functional potency compared to their more frequently dividing counterparts (Bernitz et al., 2016; Foudi et al., 2009; Qiu et al., 2014; Sacma et al., 2019; Wilson et al., 2008). Such a finding suggests that HSC potency gradually decreases with increasing rounds of cell division and that self-renewal divisions do not occur, or are rare events (Hinge et al., 2020). In line with this concept, it has previously been shown that repeated exposure to stress agonists that increase HSC division rate *in vivo*, such as inflammation and infection, can compromise HSC function (Esplin et al., 2011; Matatall et al., 2016; Pietras et al., 2014; Pietras et al., 2016; Rodriguez et al., 2009; Takizawa et al., 2017). In the setting of a murine genetic model of bone marrow (BM) failure, such challenge resulted in the exhaustion of HSC reserves and subsequent development of severe aplastic anemia, demonstrating the biological relevance of this finding (Walter et al., 2015). In this study, we sought to explore the broader implications of these findings in the context of normal hematopoiesis and specifically interrogate whether there was any evidence of regeneration of the HSC pool in this setting.

The acute challenge of mice with the toll-like receptor 3 agonist, polyinosinic:polycytidylic acid (pI:pC), leads to induction of interferon-α, transient peripheral blood (PB) cytopenias and a parallel increase in proliferation of long-term HSCs (LT-HSCs), all of which return to homeostatic levels within 4 days (Figure S1A) (Essers et al., 2009; Walter et al., 2015). To assess the effects of repeated inflammatory challenge, wild-type C57BL/6J mice were subjected to a pI:pC dose escalation regimen, with analysis of hematologic parameters performed after a four-week recovery period (Figure S1B). One to three rounds of pI:pC treatment, corresponding to 8 to 24 individual injections over the course of 8 to 24 weeks, failed to provoke any pronounced changes in PB and BM cellularity (Figures S1C-F). While the frequency and absolute number of phenotypic and transcriptionally-defined LT-HSCs in the BM of treated mice was comparable to phosphate buffered saline (PBS)-treated controls (CON), it was possible to discern a mild but significant decrease in the frequency of quiescent LT-HSCs after two or more rounds of pI:pC treatment (Figures S1G-I, S2A-D). A more detailed assessment of LT-HSC function *in vitro*, revealed that the clonogenic potential of LT-HSCs isolated from mice treated with three rounds of pI:pC (Tx^3x^) was reduced compared to those from CON mice (Figures 1A-B). There was a marked decrease in the overall proliferative potential of individual LT-HSCs from Tx^3x^ mice, which was most profoundly exhibited in clones producing multi-potent progeny, as opposed to phenotypic LT-HSCs generating bi- or uni-lineage progeny (Figures 1C, S2E). However, there was no evidence of compromised proliferative potential within more mature HSC/progenitor populations (Figure S2F). Continuous cell fate tracking experiments using video microscopy demonstrated more rapid differentiation kinetics of Tx^3x^ LT-HSCs compared to CON LT-HSCs, as well as an accelerated exit from quiescence into first cell cycle (Figures 1D-E). This may be indicative of cell fate priming towards these outcomes as a result of past exposure to inflammation. Taken together, these data suggest that while murine hematopoiesis is restored following repeated pI:pC challenge, the functional capacity of LT-HSCs may be compromised.

**Figure 1.**
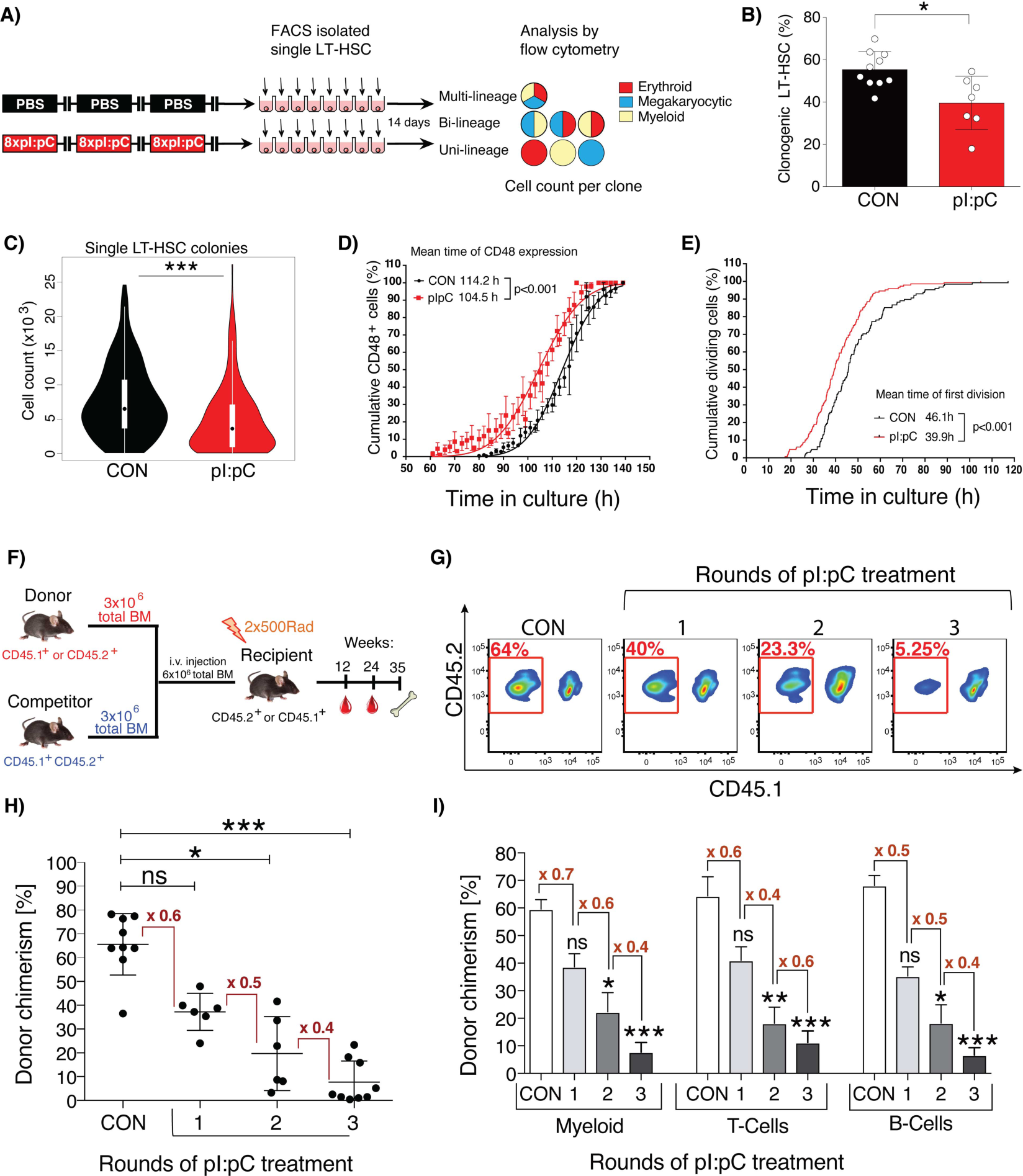
Repetitive exposure to inflammatory stress progressively compromises the functional potency of LT-HSCs. **(A)** Schematic representation of *in vitro* single cell liquid culture assay using purified LT-HSCs. **(B)** The percentage of LT-HSC clones capable of forming colonies is shown for LT-HSC isolated from CON or TX^3X^ mice (plus mean±SD, n=7-10 mice). **(C)** Violin plots representing the total number of daughter cells generated by each LT-HSC. **(D&E)** *In vitro* time-lapse microscopy-based cell tracking, evaluating: **(D)** the cumulative percentage of cells expressing CD48 versus time in culture, fitted to a Gaussian distribution curve; **(E)** the cumulative incidence of LT-HSC having undergone first cell division per unit time in culture (mean±SD, n= 128 or 148 individual LT-HSCs for CON or pI:pC groups, respectively; n=3 independent biological repeats per group). **(F-I)** Competitive repopulation assays were performed as described in methods. PB was analyzed at 24 weeks post-transplantation. **(F)** Schematic representation of the standard competitive transplantation assay **(G)** Representative flow cytometry plots of total donor leukocyte chimerism in PB. PB cells derived from donor BM isolated from pI:pC-treated or CON donors are outlined in red. **(H)** Percentage total donor leukocyte chimerism in PB for the indicated groups. Each dot represents transplantation outcome of BM from an individual treated donor mouse **(I)** Percentage donor chimerism in defined compartments of PB. Myeloid=CD11b^+^, T-cells=CD4^+^/CD8^+^, B-cells=B220^+^ (plus mean±SD, n=8-9 mice per group). ns=P>0.05, *P<0.05, **P<0.01,***P<0.001.

In order to better address the functional capacity of LT-HSCs from mice subject to repetitive induction of inflammation, competitive transplantation assays were performed using BM harvested from mice exposed to one, two or three rounds of pI:pC treatment (Figure 1F). These data demonstrated a progressive depletion of functional HSCs with increasing rounds of pI:pC challenge, correlating with an approximate two-fold reduction in the level of donor-derived hematopoiesis with each additional round of treatment (Figure 1G-I). This cumulative depletion of functional HSCs following discrete rounds of pI:pC challenge directly opposes the concept that the HSC pool is able to regenerate *in vivo* following injury, via increased self-renewal-mediated HSC expansion. In order to explore this phenomenon more comprehensively, Tx^3x^ mice were allowed to recover for 5, 10, or 20 weeks post-challenge, following which, transplantation assays were performed to ascertain if there was any evidence of regeneration of the functional HSC pool (Figures 2A, S3A-F). Surprisingly, competitive transplantation revealed that there was no significant regeneration of the reconstitution capacity of HSCs, even following an extensive recovery period (Figures 2B-C). Limiting dilution transplantation assays validated this important finding and demonstrated that there was absolutely no recovery in the absolute number of functional HSCs following pI:pC challenge, with Tx^3x^ mice still demonstrating an approximate 20-fold reduction in functional HSCs up to 20 weeks post-treatment compared to CON mice (Figures 2D, S3G-H). To address whether these findings related to compromised function of native HSCs, as opposed to defects that only become manifest upon transplantation of HSCs from treated mice, reverse transplantation experiments were performed (Figure 2E). Thus, an excess of purified HSCs harvested from non-treated wild-type mice were transplanted into Tx^3X^ recipient mice at 5, 10 or 20 weeks after treatment, in the absence of any myeloablative conditioning. In stark contrast with CON recipients, sustained multi-lineage engraftment of normal HSCs into Tx^3X^ recipients was observed, even when the transplantation was performed up to 20 weeks post-treatment (Figure 2F-G). This suggests that repeated inflammatory challenge resulted in a durable suppression of recipient HSCs, facilitating engraftment of donor HSCs for an unprecedented length of time after treatment. This data additionally demonstrates that the niche of Tx^3X^ mice is still capable of functionally supporting multilineage hematopoiesis from the non-treated donor HSCs, correlating with a lack of evidence for major permanent alterations in the composition of niche cells or spatial distribution of HSCs relative to niche landmarks following pI:pC treatment (Figures S4A-H).

**Figure 2.**
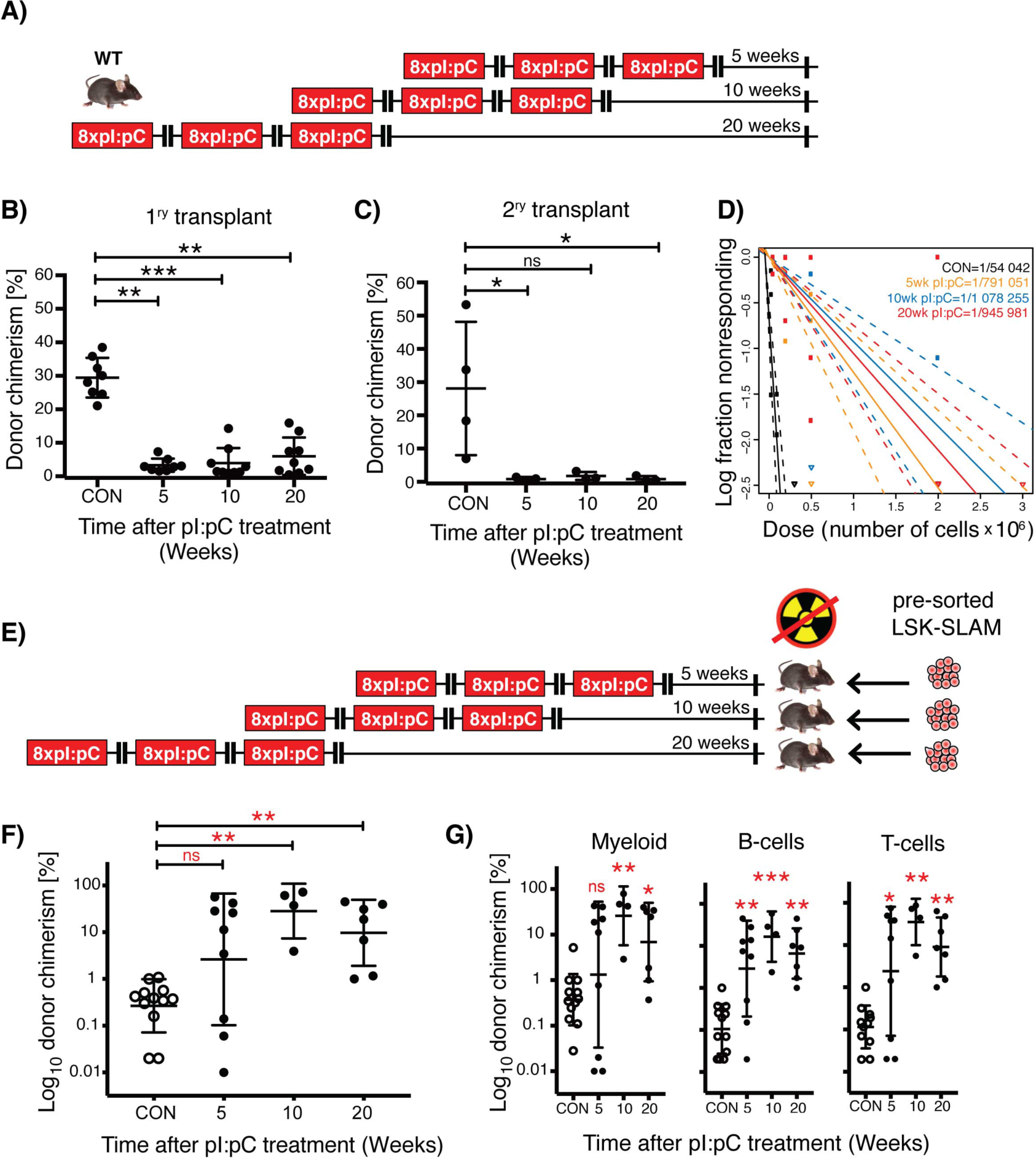
Lack of HSC functional recovery *in vivo* following inflammatory stress. **(A)** Schematic representation of treatment schedule incorporating increasing duration of recovery post-challenge with pI:pC. **(B&C)** Serial competitive repopulation assay using BM harvested from mice at indicated time points post-treatment: **(B)** Percentage total donor leukocyte chimerism at 24 weeks post-transplantation in primary recipients; **(C)** Percentage total donor leukocyte chimerism at 24 weeks post-transplantation in secondary recipients. Each dot represents transplantation outcome of BM from an individual treated mouse or primary recipient mouse (plus mean±SD). **(D)** Limiting dilution transplantation assays to determine LT-HSC frequency in BM isolated from femora of individual mice. 95% confidence intervals are indicated with dashed lines (n=6-9 recipients per dilution, per donor, representing analysis of BM from 3-4 individual treated donor mice). **(E)** Schematic representation of reverse transplant experiment. Mice exposed to the indicated treatment regimen were injected i.v. with saturating doses of purified donor HSCs in the absence of any conditioning with irradiation. **(F&G)** Percentage donor contribution in PB at 24 weeks post-reverse transplantation to the following defined populations: **(F)** total leukocytes; **(G)** Myeloid (CD11b^+^/GR-1^+^), B-cells (B220+) and T-cells (CD4^+^/CD8^+^). Each dot indicates an individual treated recipient (plus mean ± SD). ns=P>0.05, *P<0.05, **P<0.01, ***P<0.001.

To determine whether this progressive loss of HSC functionality was linked to broad systemic inhibitory effects of inflammation, or was rather associated with a cumulative increase in the *in vivo* division history of HSCs, label-retention experiments were performed using the Scl-tTA;H2B-GFP mouse model (Wilson et al., 2008), which facilitates inducible expression of GFP fused to histone H2B in LT-HSCs. Thus, one week after inducing the chase period, mice were injected with one treatment round of pI:pC or PBS (Tx^1X^) (Figure 3A). Label-retaining LT-HSCs (LRCs), which demonstrated a limited proliferative response to treatment, were then prospectively identified and isolated from their non-label-retaining (nonLRCs) counterparts based on levels of GFP fluorescence (Figure 3B). As expected, compared to PBS-treated controls, Tx^1X^ mice had fewer BM LRCs and a reduced average level of GFP fluorescence in the LT-HSC compartment, confirming that this treatment regimen enforced LT-HSC division *in vivo* (Figures 3C-D). When the functional potency of LT-HSCs was assessed *in vitro*, LRCs in Tx^1X^ group maintained their proliferative potential despite exposure to systemic inflammation (Figure 3E). This demonstrates that LRCs in Tx^1X^ mice were protected from the inhibitory effects of inflammation, and that an overall decrease in potency of LT-HSCs results from a decreased frequency of LRCs within this compartment. This suggests that inflammation-associated loss of LT-HSC potency is directly linked to proliferation history and that self-renewal divisions do not appear to take place *in vivo* under these conditions.

**Figure 3.**
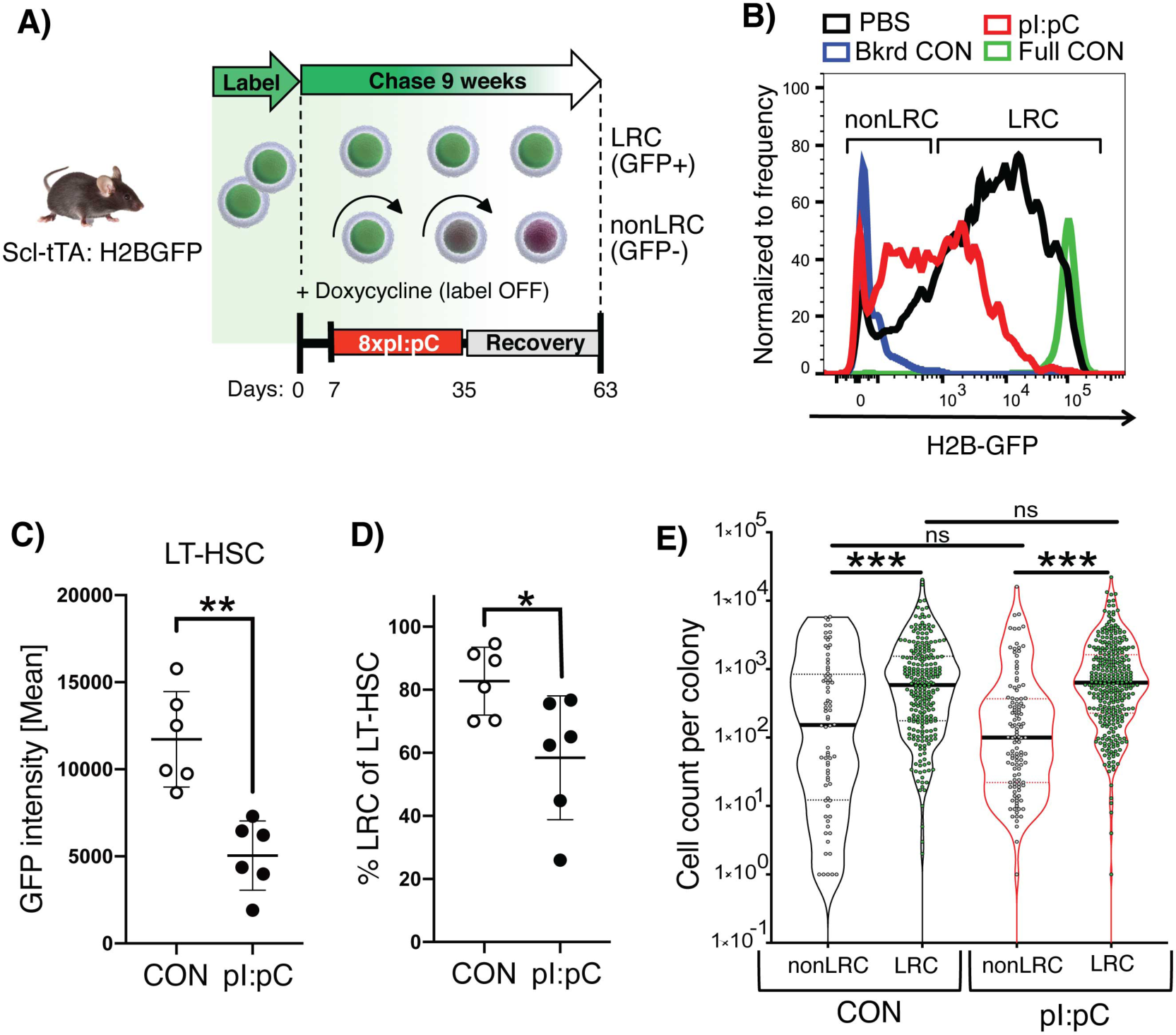
Inflammation-associated cell division leads to loss of LT-HSC potency. **(A)** Schematic representation of combined label retention and treatment schedule. Scl-tTA;H2B-GFP mice were treated with pI:pC or PBS (CON) as indicated. Label chase was induced by sustained administration of doxycycline starting 7 days before pI:pC/PBS treatment. Flow cytometry analysis/sorting was performed on BM at 8 weeks after initiation of pI:pC/PBS treatment. **(B)** Representative flow cytometry histograms of GFP fluorescence in LT-HSCs from PBS and pI:pC treated mice. Background fluorescence (Bkrd CON) and fully labeled (Full CON) controls are indicated **(C)** Mean fluorescent intensity of GFP in LT-HSCs and **(D)** the proportion of LRCs within the LT-HSC population in CON and pI:pC treated mice. Each dot represents a single mouse (plus mean±SD). **(E)** Violin plots showing the number of progeny generated per individual LRC or nonLRC following 14 days *in-vitro* culture (n=4-5 mice per group, n=280, 364, 293 or 260 analyzed clones for nonLRC CON, LRC CON, nonLRC pI:pC and LRC pI:pC, respectively. Solid lines represent median, dashed lines interquartile range, ns=P>0.05, *P<0.05, **P<0.01, ***P<0.001).

To ascertain whether this irreversible depletion of BM HSC reserves eventually results in compromised hematopoiesis, eight week old mice were subjected to eight rounds of pI:pC treatment (Tx^8X^) and were then either analyzed at 18 months of age, or were analyzed once they reached 24 months of age (Figure S5A). As previously described for Tx^3x^ mice, functional HSCs were depleted in the BM of Tx^8X^ mice, as assessed by competitive transplantation, while the PB and BM parameters were not dramatically altered relative to CON mice, when analyzed 2 months after treatment (Figures S5B-F). However, at 24 months of age, Tx^8X^ mice developed mild PB cytopenias and BM hypocellularity associated with increased BM adipocytes (Figures 4A-F), consistent with the concept that the depletion of functional HSCs was causative for the subsequent decline in output of mature hematopoietic cells.

**Figure 4.**
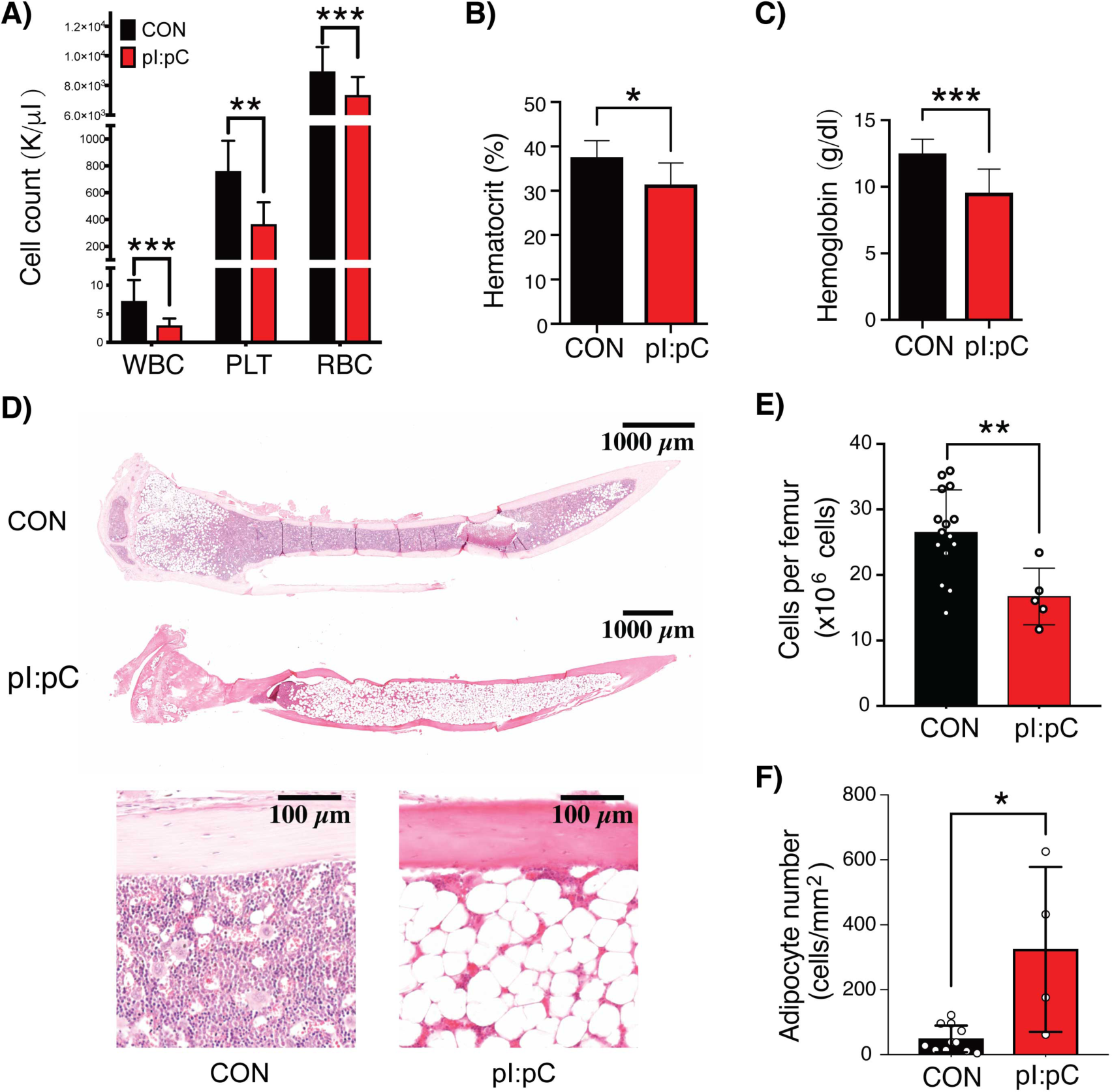
Repeated exposure to inflammatory stress provokes clinically relevant features of ageing. Mice were challenged repeatedly with pI:pC or PBS in early/mid-life as illustrated in Figure S5A. At 24 months of age, the following hematologic parameters were assessed: **(A)** Leukocyte (WBC), platelet (PLT) and red blood cell (RBC) counts in PB; **(B)** PB hematocrit; **(C)** Hemoglobin; **(D)** Representative whole mount H&E sections of tibiae; **(E)** BM cellularity per femur; and **(F)** Microscopy-based enumeration of adipocyte density within medullary cavity of tibiae (n=5-15 mice per group). Circles represent individual treated aged mice (plus mean±SD). *P<0.05, **P<0.01, ***P<0.001.

Taken together, our results demonstrate that inflammatory challenge results in a progressive irreversible depletion of functional LT-HSCs, linked to accumulating division history in the setting of little to no self-renewal. This data provides evidence that discrete episodes of exposure to inflammatory stress can have a cumulative effect on stem cell attrition over long periods of time. Such a model has clear implications for how functional HSCs can become depleted during aging.

These findings would appear to have parallels with the often cited but poorly defined phenomenon of stem cell exhaustion, where excessive self-renewal divisions, such as those induced by serial HSC transplantation, are thought to eventually lead to the functional collapse of HSC clones (Orford and Scadden, 2008). However, the label retention data that we present, rather suggests that relatively modest stimulation of the quiescent HSC pool can lead to functional attrition of the cells that respond to such stress stimuli, while cells that remain dormant are preserved. In this setting, the residual HSC and progenitor pool is clearly capable of sustaining hematopoiesis and responding to any subsequent challenge. It is only after an extended passage of time following depletion of the functional HSC pool that one observes compromised production of mature blood cells. This partly mimics the mild PB cytopenias, BM hypocellularity and accumulation of BM adipocytes, that together comprise typical features of aged human hematopoiesis, but which are not normally observed in experimental mice housed under laboratory conditions (Biino et al., 2013; Guralnik et al., 2004; Hartsock et al., 1965; Tuljapurkar et al., 2011).

The surprising revelation that LT-HSCs demonstrate little to no capacity for self-renewal in response to inflammatory challenge may be more broadly relevant to other adult stem cell populations and could explain the link between so-called inflammaging and compromised tissue maintenance and repair in the elderly.

## Acknowledgements

We thank members of the Division of Experimental Hematology for supporting the experimental work described in this manuscript, and Steven Lane, Thordur Oskarsson, Martin Sprick and Leonard Zon for critical proofreading of this work. We also thank the Center for Preclinical Research DKFZ core facility; the Flow Cytometry DKFZ core facility; the Single Cell Open Lab DKFZ Core Facility; and Damir Krunic from the Light Microscopy DKFZ core facility. This work was supported by funding from the German Research Foundation (DFG) SFB873 (MDM and MAGE), FOR2674 (MDM, DBL, BB, KR and JPM) and SFB834 (MAR and MF); the Deutsche Jose Carreras Leukämiestiftung (grant R15/09 to MDM and 10R/2017 to MAR); the Fritz Thyssen Stiftung (grant 10.16.1.023MN to MDM); the Helmholtz Zukunftsthema Aging and Metabolic Programming (AMPro) ZT-0026 (MDM and DBL); the DKFZ-MOST German-Israel Cooperative Research Program (MDM); the Cancer Transitional Research And Exchange Program (Cancer-TRAX) within the German-Israeli Helmholtz International Research School (SS); the National Institutes of Health RO1 DK056638 and R01 DK112976 (PF); the Wilhelm-Sander Foundation (grant 2018-116.1 to MAR); and the Dietmar Hopp Stiftung (MDM and MAGE).

## Author contributions

RB, BB, SH, DBL, MAGE, KR, PSF, MAR and MDM designed and directed the experimental scheme of work; RB, PK, MF, AMM, MBS, SVP, NA, BM, JK, SS, DW, AP, MJCG, VW and JPM performed experiments; RB, PK, MF, FF, FB, BB, SH, DBL, MAGE, THL, JPM, KR, PSF, MAR and MDM carried out data analysis and/or interpretation of experimental data; THL and RB performed statistical analysis of the data; RB, MF, NA, FF, FB, PSF, MAR and MDM wrote the manuscript.

## Additional information

Correspondence and requests for materials should be addressed to Michael D. Milsom.

## Methods

### Animals and animal experiments

All animal experiments were approved by the local Animal Care and Use Committees of the “Regierungspräsidium Karlsruhe für Tierschutz und Arzneimittelüberwachung”. Mice were maintained under specific pathogen-free conditions in individually ventilated cages at the German Cancer Research Center (DKFZ, Heidelberg) animal facility. Wild-type mice (C57BL/6 or B6.SJL-Ptprca Pepcb/BoyJ) were obtained from Harlan Laboratories, Charles River Laboratories, or Janvier Laboratories. Unless otherwise indicated, mice were 8 to 16 weeks old when experiments were initiated. H2B-GFP and ScltTA mice have been previously described (Bockamp et al., 2006; Tumbar et al., 2004; Wilson et al., 2008). H2B-GFP and ScltTA mice on a C57BL/6 background were crossed in order to perform a label retaining cell (LRC) assay. For the LRC assays, doxycyline treatment was performed by supplementing the drinking water of experimental mice with 2 mg/ml doxycyline citrate (Sigma), sweetened with 20 mg/ml sucrose. Doxycycline-supplemented drinking water was sustained for the duration of the label chase period.

### Treatment with pI:pC

To mimic a repetitive sterile inflammatory response *in vivo*, mice were serially injected intraperitoneally (i.p.) with 5 mg/kg high molecular weight polyinosinic:polycytidylic acid (pI:pC, InvivoGen), which had been reconstituted in sterile physiologic saline precisely as described in the manufacturer’s instructions. Control mice were injected with the same volume of PBS (Sigma Aldrich).

### Bleeding and bone marrow isolation

Peripheral blood (PB) was collected by puncturing the craniofacial capillary bed of the mice and up to 100 µl of PB was collected into EDTA-coated tubes. PB counts were evaluated using a Hemavet 950 FS veterinary blood cell counting machine (Drew Scientific).

For the purification of murine bone marrow (BM) cells, hind legs (femora, tibia and iliac crests) and vertebrae were dissected by removing adherent soft tissue and the spinal cord using a scalpel. Bones were either crushed or flushed, and the resulting cell suspension was filtered through 40 µm cell strainers (Greiner Bio-One) and re-suspended in ice cold 2 % (v/v) FCS/PBS (PAA Laboratories/Sigma Aldrich) following centrifugation. When BM cells were pooled from multiple mice, bones were gently crushed in Iscove’s modified Dulbecco’s medium (IMDM, Life Technologies). Bones from individual mice were harvested by flushing the cells out of two femurs into 2 ml ice cold 2 % (v/v) FCS/PBS or PBS using a 1 ml syringe fitted with a 23 gauge needle.

### Competitive repopulation transplantation assays

CD45.2 or CD45.1 C57BL/6 recipient mice were subject to total body irradiation (2 x 5 Gy TBI, Bestrahlungsgerät/Buchler GmbH, caesium source) 2 to 16 hours prior to BM transplantation. Recipients were co-injected intravenously (i.v.) with a mixture of 3 x 10^6^ WT CD45.1/CD45.2 whole BM competitor cells and 3 x 10^6^ WT CD45.1 or CD45.2 whole BM test donor cells (Figure 1F). BM from each individual test mouse (pI:pC or PBS) were separately transplanted into individual recipient mice. However, in order to facilitate high reproducibly, the co-injected competitor BM cells came from a common pool of cells, isolated from at least two donors. The same pool of competitor BM was used for all transplanted mice within each experimental repeat. To evaluate the repopulation potential of the test BM populations, PB and BM of recipient mice were analyzed by flow cytometry at three, six and eight months post-transplantation. PB and BM were stained with monoclonal antibodies against CD45.1 and CD45.2, in order to discriminate between reconstitution from test donor cells; competitor donor cells; and endogenous recipient cells. PB and BM were additionally stained with antibodies directed against B220, CD4, CD8a, CD11b; CD5, Gr-1, Ter-119, c-Kit, Sca-1, CD150 and CD48, in order to evaluate donor reconstitution within defined mature and immature cell populations. An aliquot of the input cell mix was separately stained and evaluated by flow cytometry to validate the correct ratio of cells was injected into recipient mice.

### Limiting dilution transplantation assays

Total BM cells from pI:pC or PBS treated CD45.1 C57BL/6 mice were injected i.v. at different cell doses into cohorts of lethally irradiated CD45.2 recipient mice (2 x 5 Gy TBI), together with a fixed rescue dose of 2 x 10^5^ CD45.1/CD45.2 BM cells. The BM of each test mouse was transplanted into 24 individual recipient mice. Recipients of BM from pI:pC-treated mice were injected with either 3 x 10^6^ (6 recipients), 2 x 10^6^ (60 recipients), 5 x 10^5^ (60 recipients), 2 x 10^5^ (60 recipients) or 5 x 10^4^ BM cells (60 recipients), while recipients of PBS-treated BM received either 3 x 10^5^ (18 recipients), 1 x 10^5^ (25 recipients) or 3 x 10^4^ BM cells (25 recipients). At least three different donor mice from each experimental group were individually assessed using this methodology. Engraftment was assessed by flow cytometry analysis of PB at six months post-transplantation. Mice that demonstrated ≥ 1% donor-derived contribution to both myeloid (Gr-1^+^ and/or CD11b^+^) and lymphoid (B220^+^ and/or CD4^+^ and/or CD8^+^) lineages in the PB were scored as positive (responding) for engraftment. To estimate the frequency of repopulating HSCs in the BM, a limiting dilution calculation was performed using the web-based ELDA software provided at http://bioinf.wehi.edu.au/software/elda/ (Hu and Smyth, 2009), using the number of responding mice at each cell dose as input data.

### Reverse transplantation experiments into non-conditioned mice

Recipient CD45.1 mice were treated with three rounds of pI:pC or PBS, as illustrated in Fig. 2e. At 5, 10 or 20 weeks after treatment, mice were injected i.v. with saturating doses of (1.5 - 3 x 10^3^) FACS-purified Lineage^-^, Sca-1^+^, c-Kit^+^, CD150^+^ BM cells isolated from a CD45.2 donor. Importantly, the recipients were not subject to any additional myelosuppressive conditioning, such as total body irradiation or chemotherapy. The level of donor chimerism in defined cell populations of the PB and BM was assessed at six and eight months post-transplantation, respectively.

### Flow Cytometry Analysis and Sorting

#### Fluorescent staining of PB and BM

PB and BM were stained with monoclonal antibodies directed against specific cell surface epitopes as detailed in Table S1. All antibodies had previously been titrated and were used at a concentration where the mean fluorescent intensity plateaus. Cells were incubated with the antibody mix in 2 % v/v FCS/PBS for 30 min. at 4 °C, washed with 2 % FCS/PBS and then re-suspended in 2 % FCS/PBS containing 7-amino actinomycin (7-AAD, Invitrogen) at a concentration of 5 µg/ml. For PB samples, an additional erythrocyte lysis step with 1 ml ACK lysing buffer (Lonza) for 10 min. at room temperature was carried out after the staining.

#### Flow cytometry analysis

After surface staining, cells were analyzed by flow cytometry using either an LSRII or an LSRFortessa cytometer (Becton Dickinson) equipped with 350 nm, 405 nm, 488 nm, 561 nm and 641 nm excitation lasers. Prior to the analysis of cells, compensation was manually adjusted using OneComp eBeads (eBioscience) stained with single antibodies. Analysis of flow cytometric data was performed using FlowJo software (Tree Star). If not indicated otherwise, populations were gated according to the markers listed in Supplementary Information.

#### Cell cycle analysis

BM cells were stained with the LT-HSC antibody panel (Supplementary Information). After surface staining, cells were lysed using ACK lysing buffer, washed with PBS and fixed with BD Cytofix/Cytoperm (BD Bioscience) for 20 min. at 4 °C. Then, cells were washed twice with PermWash (BD Bioscience); re-suspended in 100 µl PermWash, containing mouse anti-human Ki-67 (Supplementary Information) and incubated overnight at 4 °C. Shortly before flow cytometry analysis, the cells were incubated with Hoechst33342 in a 1/400 dilution for 10 min. at 4 °C.

#### Isolation of murine LSK/LT-HSC cells via FACS

To purify low-density mononuclear cells (LDMNCs) from BM cells, three rounds of density gradient centrifugation using Histopaque 1083 (Sigma-Aldrich) were performed at room temperature. An equal volume of BM cell suspension (2-10x10^7^ cells/ml) was carefully layered on top of an equal volume of the Histopaque 1083 in a 15 ml falcon tube (Greiner; Sarstedt). After centrifugation at room temperature at 300 g for 20 min. with the brake switched off, the LDMNC fraction was collected without disturbing the pellet. The pellet was re-suspended and re-applied to Histopaque 1083 for the second round of density gradient centrifugation. In total, three rounds of centrifugation were performed. The fractions containing the LDMNCs were pooled and washed with ice-cold PBS. For lineage depletion, the LDMNC fraction was incubated with a panel of rat anti-mouse biotin-conjugated lineage markers (4.2 µg/ml CD5, 4.2 µg/ml CD8a, 2.4 µg/ml CD11b, 2.8 µg/ml B220, 2.4 µg/ml Gr-1, 2.6 µg/ml Ter-119) for 45 min. at 4 °C. After washing with ice-cold PBS, the labeled LDMNCs were incubated with Biotin Binder Dynabeads at a ratio of 4 beads per input cell (Life Technologies) and the lineage-positive cells were depleted using a magnetic particle concentrator according to the manufacturer’s instructions (Dynal MPC-6, Invitrogen). To isolate the LSK/LT-HSC fraction by FACS, the resulting lineage-depleted cells were subsequently stained with a panel of antibodies (c-Kit, Sca-1, CD150, CD48, CD34), as indicated in Supplementary Information.

In order to maximize the yield of LRC and nonLRC LT-HSCs from ScltTA;H2BGFP mice, density gradient centrifugation was omitted and lineage depletion was performed directly after BM isolation. LRC and nonLRC LT-HSC from ScltTA;H2B-GFP on doxycycline treatment were defined as follows. Maximum GFP intensity was determined by flow cytometry of BM isolated from a control ScltTA;H2B-GFP mouse not exposed to doxycycline treatment (Full CON). The non-specific background GFP signal (Bkrd CON) was defined in LT-HSCs from H2B-GFP mice. LT-HSCs showing GFP intensity above the Bkrd CON were defined as LRCs, where LT-HSC with an overlapping GFP intensity to the Bkrd CON were defined as nonLRC, as shown in Figure 3B.

Sorting experiments were performed using a BD FacsAria I, II or III flow cytometer (BD Bioscience) at the DKFZ Flow Cytometry Service Unit, using a 100 µm nozzle and a maximum sort rate of 3 thousand cells per second. Single cell sorts directly into 96 multi-well plates were performed using the single cell precision mode, where the drop trajectory was adjusted for a 96-well plate before each sort.

### Microscopy analysis

#### Immunofluorescence imaging of BM niche components

Whole mount staining of HSCs in sternum bone marrow was performed as previously described (Kunisaki et al., 2013). Briefly, Alexa Fluor 647-anti-CD144 (BV13) and Alexa Fluor 647-anti-CD31 (MEC13.3) (from Biolegend) were injected i.v. 10 minutes before euthanizing mice, in order to stain BM endothelial cells in vivo. Sternal bones were collected and transected with a surgical blade into 2-3 fragments. The fragments were bisected sagitally for the BM cavity to be exposed, and then fixed with 4 % PFA for 30 min. After rinsing with PBS, bone pieces were blocked/permeabilized in PBS containing 20 % (v/v) normal goat serum and 0.5 % (v/v) TritonX-100. Primary antibodies were incubated for approximately 36 hours at room temperature. After rinsing the tissue with PBS, the tissues were incubated with secondary antibodies for 2h. The primary antibodies used were biotin-anti-Lineage (TER119, RB6-8C5, RA3-6B2, M1/70, 145-2C11) (from BD Biosciences); biotin-anti-CD48 (HM48-1), biotin-anti-CD41 (MWReg30) (from eBioscience); and Alexa Fluor 647-anti-CD144 (BV13), PE-anti-CD150 (TC15-12F12.2) (from Biolegend). The secondary antibody used was Streptavidin eFluor 450 (eBioscience). Images were acquired using ZEISS AXIO examiner D1 microscope (ZEISS) with a confocal scanner unit, CSUX1CU (Yokogawa), and reconstructed in three dimensions with Slide Book software (Intelligent Imaging Innovations). Two-sample Kolmogorov-Smirnov tests were used for comparisons of distribution patterns. Statistical analyses were performed using GraphPad Prism 6 software.

#### Histology, hematoxylin and eosin (H&E) staining

Tibiae were fixed in 10 % formalin in PBS (v/v) for not longer than one week and decalcified for five days in 0.5 M EDTA (Ethylenediaminetetraacetic acid) buffer (pH 7.2). Bones were dehydrated in the Tissue-TeK-VIP Sakura tissue processor overnight and subsequently paraffin embedded using the HistoStar embedding workstation (Thermo Scientific). Embedded bones were cut with a Microtome (Microm HM 355S, Thermo Scientific) and stained with Hematoxylin/Eosin (H&E). In brief, bone sections were de-paraffinized and rehydrated: 3 times in xylol for 5 minutes, 2 times in 100 % ethanol, 2 times in 96 % ethanol, 1 time in 70 % ethanol and lastly transferred to VE water. Slides were then stained in Mayer’s haematoxylin for 5 minutes and then rinsed under running tap water for 5 minutes. Subsequently, they were dipped into acid EtOH (0,25 % (v/v) HCl in 70 % (v/v) EtOH) and washed until the sections were stained blue. They were counterstained with eosin for 1 minute, dipped into 95 % EtOH and 100 % EtOH and put into xylene for 15 minutes. The sections were then embedded in mounting media and dried overnight. Imaging was performed with a Zeiss Axioplan widefield microscope.

#### Adipocyte quantification in H&E sections

Adipocytes were counted from H&E stained tibia sections. The images were taken with an Axio Plan Zeiss Microscope equipped with Axio Cam ICc3 Zeiss (2.5x magnification) and processed with the ZEN program 2011. Adipocyte quantification was performed in Fiji (ImageJ), where individual bone marrow adipocytes, as defined by the following parameters: size (40-2000 pixel) and shape (circularity 0.4-1.00), were counted in a predefined surface area.

### Single cell transcriptomic analysis

#### Single cell RNA-sequencing (scRNAseq)

LT-HSCs were purified by FACS, and single LT-HSC cells were subsequently captured on a small sized IFC using the Fluidigm C1 system. Briefly, cells were washed, re-suspended in PBS supplemented with C1 suspension buffer in a 4:1 ratio and 400 cells/µl were loaded onto the chip. After cell capture, each position on the chip was imaged and only single cells were included in the downstream library preparation and analysis. cDNA was then amplified with the SMARTer Ultra Low RNA kit (Clontech) including ERCC RNA spike-ins (ThermoFisher Scientific# 4456740). Bulk controls were also processed for each C1 run using 100 cells and the same reagent mixes as used for the C1.

Amplified cDNA was checked with the TapeStation to assess both quality and yield. Sequencing libraries were produced with the Illumina Nextera XT kit according to the adopted Fluidigm protocol. All single cells from one C1 run (about 70 cells on average) were pooled and sequenced 1x50 bp reads on an Illumina HiSeq 2000 machine resulting in 2-3 million reads per cell.

#### scRNAseq bioinformatic analysis

For each cell, reads were aligned to the murine genome (ERCC sequences concatenated to GRCm38.p4 version 84, softmasked) with STAR version 2.5. For each cell, between 70 and 90 % of the reads were uniquely mapped. Raw counts were quantified from position-sorted alignment files with HTSeq-count using mode ’union’ and default quality thresholds of 10. Cells were excluded as low quality if more than 40 % of counts were in ERCCs, or if the counts in murine exons were more than 10 % mitochondrial or less than 0.5 Mio in total. In addition, cells for which less than 2,000 genes were expressed were excluded; resulting in a total of 564 cells passing quality control. Size-factor normalization (Love et al., 2014) was used to identify variable genes using a log-linear fit capturing the relationship between mean and squared coefficient of variation (CV) of the log-transformed, TPM data (Brennecke et al., 2013). Genes with a squared CV greater than the estimated squared baseline CV were then considered as variable beyond technical noise. This filter for highly variable genes resulted in 5176 genes. This set of variable genes was used as input for downstream analysis, including visualization and clustering.

A projection analysis was performed to integrate our own data with a larger hematopoietic dataset covering a wider range of blood stem and progenitor cells (Nestorowa et al., 2016). To this end, the intersection of variable genes identified in (Nestorowa et al., 2016) and our data was established. A diffusion map representation of the 1656 cells from (Nestorowa et al., 2016) was then generated, based on the 1616 genes that were variable above technical noise both in our data and the data from (Nestorowa et al., 2016). Our cells were then projected into the diffusion map span based on the diverse set of stem and progenitor cells from (Nestorowa et al., 2016) using the destiny R package (Angerer et al., 2016).

### *In vitro* single cell growth assays

#### Single cell LT-HSC clonogenic assay

LT-HSCs were directly flow sorted as individual cells per well into retronectin pre-coated ultra-low attachment 96-well plates (Sigma-Aldrich) in serum-free medium (StemSpan SFEM) containing 1 % (v/v) penicillin/streptomycin, 1 % (v/v) L-glutamine, and recombinant murine cytokines that facilitate HSC growth and *in-vitro* differentiation into erythroid, myeloid and megakaryocytic lineages (10 ng/ml Flt3-Ligand, 50 ng/ml SCF, 10 ng/ml TPO, 5 ng/ml IL-3, 10 ng/ml IL-11, 0.3 IU/ml Epo, 20 ng/ml IL-7, all from PeproTech). During the clonogenic expansion, the single LT-HSCs were cultured under hypoxic conditions (5 % O_2_), 37 °C, 5 % CO_2_ for 12-14 days. The differentiation potential (myeloid, erythroid, megakaryocytic) and proliferative capacity (relative number of cells per colony) for each LT-HSC colony was enumerated by flow cytometry, essentially as previously described (Haas et al., 2015). Briefly, cells were directly stained with antibodies in the well and the entire content of the well was run through the flow cytometer. The percentage of cells contributing to the myeloid, erythroid or megakaryocytic lineages within each LT-HSC colony was determined by the expression of lineage specific markers (Gr-1/CD11b, Ter-119, CD71, CD4, CD42d).

#### Classification of the differentiation potential of single cell derived LT-HSC clones

LT-HSCs were classified into 7 subgroups depending on whether they had the potential to differentiate into a single cell type (unipotent cells: myeloid, erythroid or megakaryocytic), into two cell types (bipotent cells: myeloid-erythroid, myeloid-megakaryocytic, megakaryocytic-erythroid) or into all three (multipotent cells). Cells were ascribed to these groups as follows. In order to account for different base frequencies of the three descendent cell types all observed cell numbers were first normalized by the maximum number of the respective cell type over all observed cells. Next, proportions of the three normalized cell counts were calculated for each colony. In theory, each unipotent LT-HSC should produce 100 % of the specified descendent cells, bipotent LT-HSC should ideally produce close to 50 % each of the normalized numbers of descendants and multipotent cells should produce 33.3 % each. Thus, we plotted these theoretical subgroup means as well as the actual cells in a graph illustrating the descendent proportions. As the proportions add up to 100 %, a two-dimensional plot of any two of the proportions is sufficient for this analysis. Each of the actual cells was then classified into the subgroup it was closest to based on Euclidian distance. For all data sets, each of the cells could thus be classified into one subgroup and the proportion of the 7 subgroups could be determined. Differences in the distribution of these proportions where then tested for statistical significance using a Chi-Square test, using a significance level of alpha=5 %. All calculations were performed using R, Version 3.2.0.

### Statistical analysis

Unless otherwise indicated, data are presented as mean +/- standard deviation. Statistical analyses were carried out in comparison to the control group. For pairwise comparisons, two-sided unpaired non-parametric t-tests were applied (Fig.1B, C; Fig.3 C, D; Fig.4 A, B, C, E, F). Comparisons of more than two groups were performed by one-way analysis of variance (ANOVA) on ranks. If the ANOVA provided evidence that group means differed, Dunn’s multiple comparison tests were applied to determine which means amongst the set of means differed from the rest (Fig.1H, I; Fig.2B, C, F, G; Fig.3E). To evaluate the cumulative frequency distribution of CD48 expression on LT-HSCs, the data was fitted to Gaussian distribution curves by least squares regression. Best-fit values of the control and treatment datasets were compared to each other by extra sum-of-squares F test (Fig.1D). Log-rank Mantel-Cox test was used to test for statistically significant changes of first LT-HSC division between the control and treatment group (Fig.1E). Variables that showed skewed distribution were Log10 transformed (Fig.2F, G). Statistical significance is indicated by one (P < 0.05), two (P < 0.01) or three (P < 0.001) asterisks. Analyses were performed using GraphPad Prism 5.0b software (GraphPad Software, Inc., SanDiego, CA, http://www.graphpad.com).

### Time-lapse imaging and single cell tracking

Time-lapse imaging and cell tracking were performed as previously described (Cabezas-Wallscheid et al., 2017; Haetscher et al., 2015; Rieger et al., 2009). LT-HSCs were FACS purified from pI:pC-treated or control mice and seeded in 24-well plates equipped with silicon culture inserts (IBIDI, Martinsried, Germany). Cells were pre-cultured in StemSpan SFEM medium (StemCell Technologies) supplemented with 10 ng/ml Flt3-Ligand, 50 ng/ml SCF, 10 ng/ml TPO, 5 ng/ml IL-3, 10 ng/ml IL-11, 0.3 IU/ml Epo, 20 ng/ml IL-7 (PeproTech) recombinant murine cytokines and 0.1 ng/ml rat anti-mouse CD48-PE (clone HM48-1, eBioscience) for 17 h in a standard cell culture incubator at 37 °C and 5 % CO2 for CO2 saturation, before being gastight sealed with adhesive tape for live-cell microscopy. Time-lapse imaging was performed using a CellObserver system (Zeiss, Hallbergmoos, Germany) at 37 °C. Phase contrast images were acquired every 2-3 min over 7 days using a 10x phase contrast objective (Zeiss), and an AxioCamHRm camera (at 1388x1040 pixel resolution) with a self-written VBA module remote controlling Zeiss AxioVision 4.8 software. PE fluorescence (Filter set F4-004, AHF Analyzetechnik at 600ms) was detected every 2 hours. Cells were individually tracked for their fates (apoptosis, division, loss of stemness) using a self-written computer program (TTT) in concert with manual verification and analysis of results. The generation time of an individual cell was defined as the time span from cytokinesis of its mother cell division to its own division. Dead cells were identified by their shrunken, non-refracting and immobile appearance. Induction of differentiation was detected by the appearance of PE fluorescence (CD48 expression). The analysis did not rely on data generated by an unsupervised computer algorithm for automated tracking.

---

**Figure S1.**
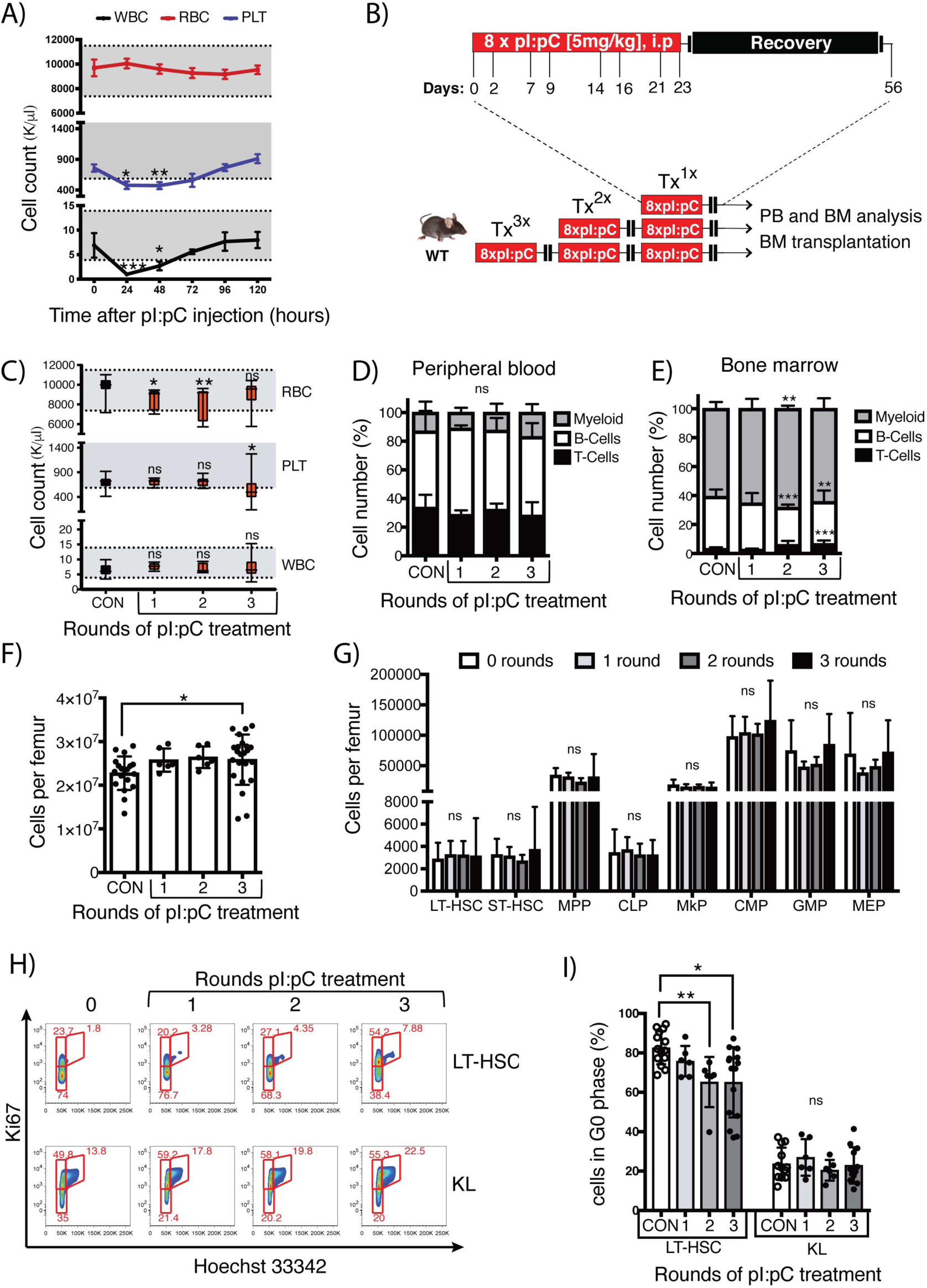
Repeated pI:pC treatment has a negligible sustained impact upon the quantitative and qualitative composition of peripheral blood and bone marrow. **(A)** Time course analysis of peripheral blood (PB) cell counts following a single injection of C57BL/6J mice with pI:pC. The normal range values for white blood cell (WBC), red blood cell (RBC) and platelet (PLT) counts are indicated in grey. The mean ± SD are indicated. n=6-9 mice per time point. **(B)** Schematic representation of pI:pC dose escalation regimen. Each round of treatment (TX^1X^) consists of 8 i.p. injections with pI:pC on the indicated days, followed by a 33 day recovery period prior to the next round of treatment. **(C)** PB cell counts of PBS-treated control mice (CON) and mice treated with 1, 2 or 3 rounds with pI:pC, as indicated in Figure S1B. PB counts were evaluated 5 weeks after the last pI:pC injection, n=5-29 per group. Box and whiskers plots indicate median, interquartile range and minimum to maximum values. **(D-E)** The relative frequency of myeloid (CD11b^+^/GR-1^+^), B- (B220^+^) and T-cells (CD4^+^/CD8^+^) was measured by flow cytometry in PB **(D)** and BM **(E)** of CON and pI:pC treated mice, n=6-18 per group. Mean±SD are shown. **(F)** BM femur cellularity was enumerated 5 weeks after the indicated treatment regimen. Mean±SD are shown. n=6-23 mice. **(G)** The relative frequency of defined BM HSC and progenitor populations, as specified in table S2, was determined by flow cytometry. Absolute frequencies were calculated by adjusting for femur cellularity for each individual mouse. n=6-12 per group. Mean±SD are shown. **(H)** Representative flow cytometry plots showing determination of cell cycle status of LT-HSC and KL progenitor cells using Hoechst and Ki67 staining. The gating strategy used to segregate cells in the G0 (Ki67 and Hoechst low); G1 (Ki67 high, Hoechst low) and S/G2/M (Ki67 and Hoechst high) phases of the cell cycle are shown, as are the percentage of LT-HSCs or KL cells within each of these gates. **(I)** Relative frequency of quiescent (G0) LT-HSC and KL cells at 5 weeks after the indicated treatment regimen. Mean±SD are shown. n=6-15 mice. Statistical significance between control and treatment groups was evaluated by one-way ANOVA on ranks with Dunn’s multiple comparison tests (*P<0.05, **P<0.01, ***P<0.001).

**Figure S2.**
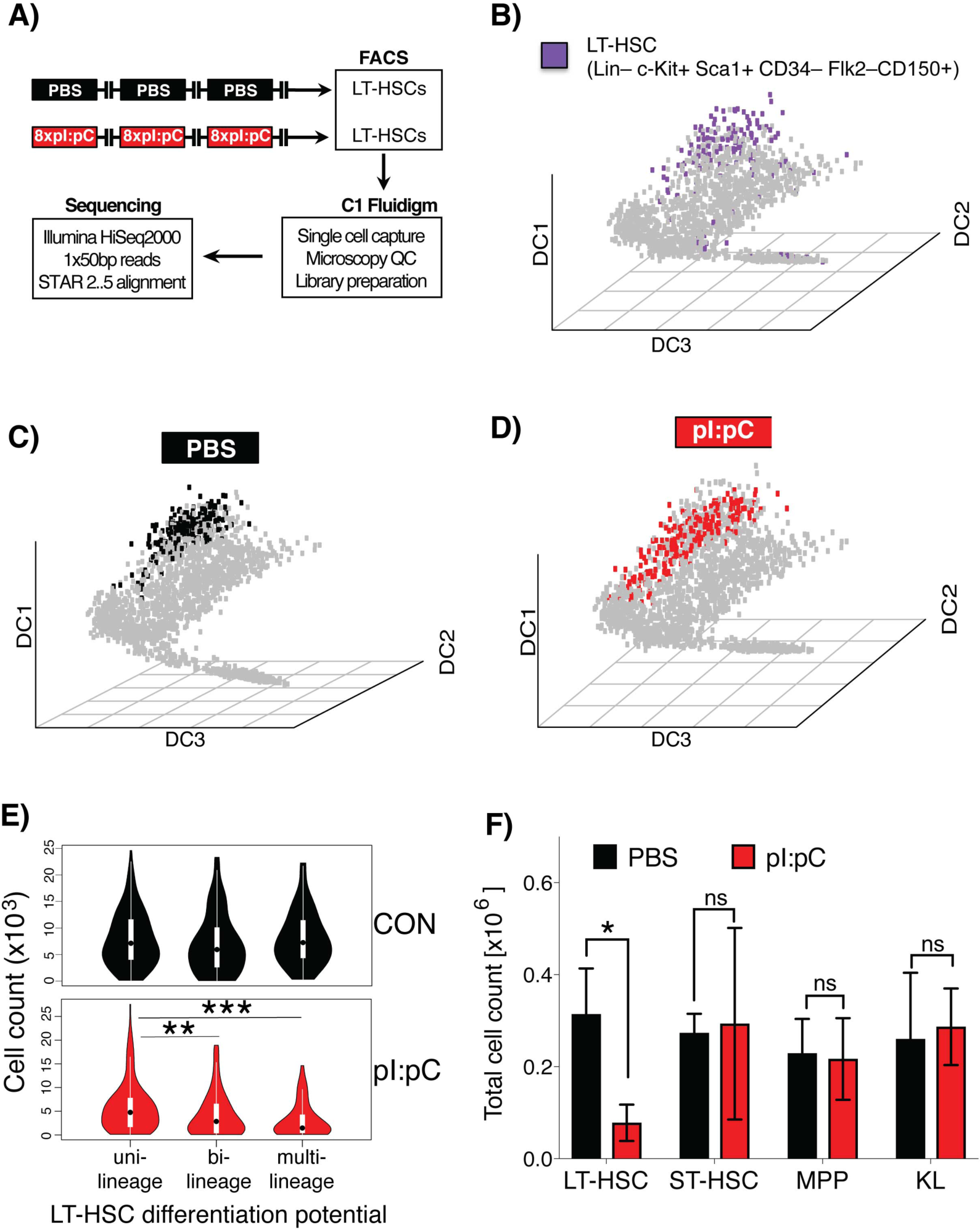
Immunophenotypic LT-HSCs from pI:pC treated mice retain a similar transcriptional identity to LT-HSCs from CON mice, but show functional impairment in *in-vitro* assays. **(A)** Schematic representation of the scRNAseq workflow. LT-HSC from CON or TX^3X^ mice were isolated by flow cytometry at 5 weeks post-treatment, and were subsequently subjected to scRNAseq using the C1 Fluidigm platform. Downstream bioinformatics analysis was performed as described in the materials and methods section. **(B-D)** scRNAseq data was comparatively analyzed relative to a publicly available set of scRNAseq data encompassing murine bone marrow HSC and progenitor cells (Nestorowa et al., 2016). A multi-dimensional diffusion map is presented with the entire HSC and progenitor compartment indicated by grey dots. **(B)** The location of immuno-phenotypically defined LT-HSCs, as previously defined in (Nestorowa et al., 2016), is indicated by purple dots in the diffusion plot. **(C-D)** The intersecting variable genes between the previously published data set and the current scRNAseq data were used to project LT-HSC from the current study onto the pre-existing data set. LT-HSC isolated from **(C)** CON or **(D)** TX^3X^ mice are indicated with black or red dots, respectively. These two populations predominantly retain the same transcriptional signatures, suggesting that the cell surface marker combinations used to identify and purify LT-HSCs, essentially mark the same cell population regardless of treatment regimen. **(E)** Violin plots representing the number of daughter cells per LT-HSC, segregated by colony type from the *in vitro* single cell liquid culture assay using purified LT-HSCs described in Figure 1A (CON n=281 and pI:pC n=272 clones; n=3 mice per group; white rectangle=interquartile range, black dot=median). **(F)** LT-HSC (200 cells); ST-HSC (200 cells); MPP (1000 cells); or KL (2000 cells) were purified from CON or TX^3X^ mice and directly placed into liquid culture. The total number of progeny was enumerated on day 10 of culture, demonstrating that the proliferative potential of LT-HSCs was compromised, while that of ST-HSCs and progenitors was not. Statistical analysis was performed using an unpaired two-tailed Student’s t-test, n=3-5 mice, data represent mean±SD. Unless otherwise stated, statistical nce between control and treatment groups was evaluated by one-way on ranks with Dunn’s multiple comparison tests (ns=P>0.05, *P<0.05, ***P<0.001).

**Figure S3.**
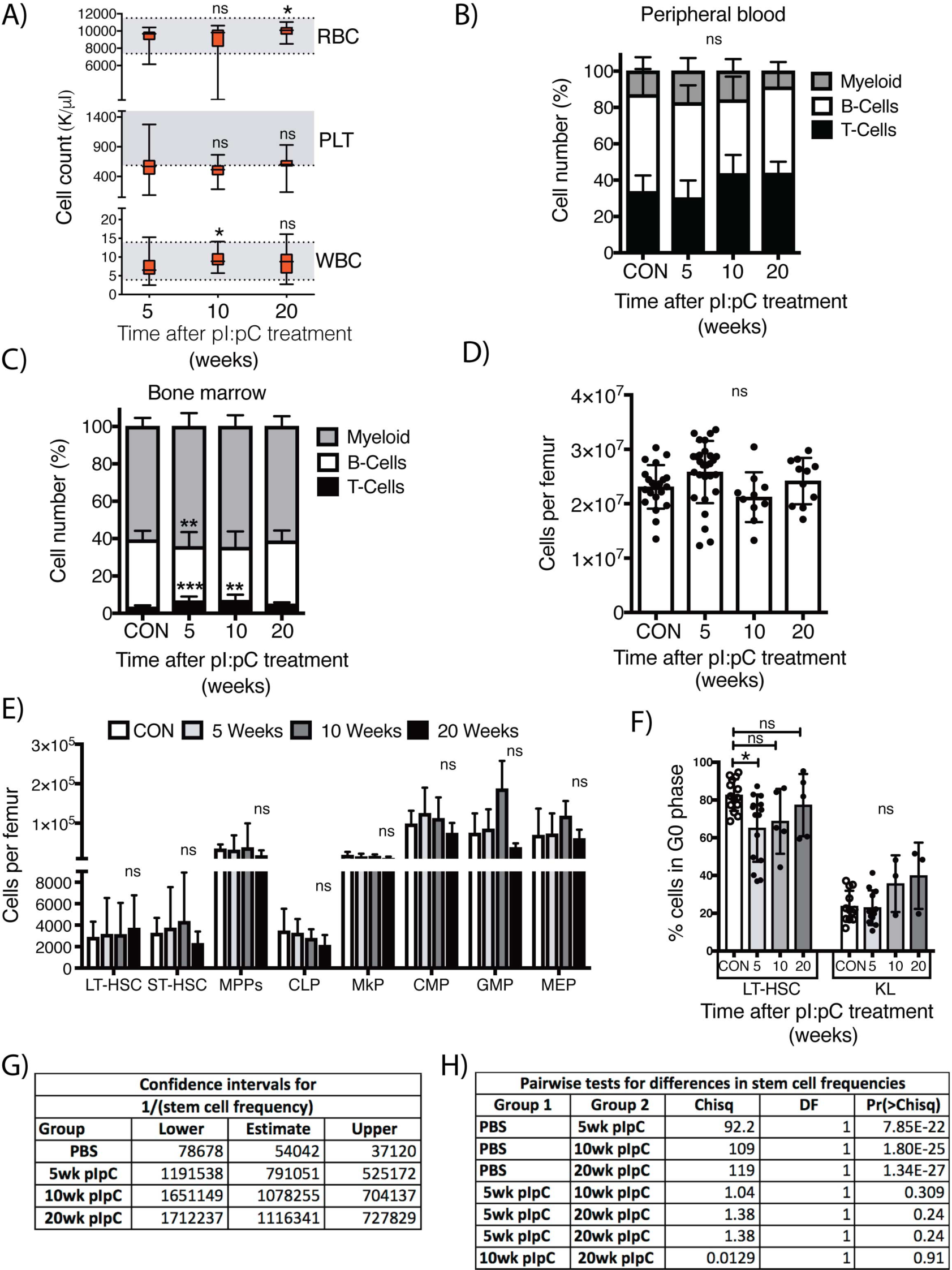
Characterization of hematological parameters and limiting dilution transplantation analysis after 5, 10 or 20 weeks recovery in TX^3X^ WT mice. **(A)** The PB counts of TX^3X^ mice was measured at 5, 10 or 20 weeks after the last pI:pC injection, Box and whiskers plots indicate median, interquartile range and minimum to maximum values, n=17-44. **(B-C)** The relative frequency of myeloid (CD11b^+^/Gr-1^+^), B- (B220^+^) and T-cells (CD4^+^/CD8^+^) was measured by flow cytometry in both the **(B)** PB and **(C)** BM of CON and TX^3X^ treated mice, at the indicated time points after treatment. Mean ± SD are shown. n=6-18. **(D)** The BM cellularity within the femora of CON and TX^3X^ mice was evaluated in mice that were sacrificed at the indicated time point after treatment. Mean ± SD are shown. n=10-28 mice. **(E)** The absolute number of cells within defined immunophenotypic compartments (see Table S2) within the BM of CON and TX^3X^ mice at the indicated time point post-treatment was assessed by flow cytometry. This was followed by adjustment to take account of BM cellularity for each individual mouse, n=6-12. **(F)** The percentage of quiescent (G0) LT-HSC and KL cells within the BM of CON and TX^3X^ mice, harvested at the indicated time point after treatment, is shown. Mean ± SD, n=3-15 mice. Statistical significance between control and treatment groups was evaluated by one-way ANOVA on ranks with Dunn’s multiple comparison tests (ns=P>0.05, *P<0.05, **P<0.01, ***P<0.001). **(G-H)** The statistical analysis of the limiting dilution experiment shown in Figure 2D is shown in these tables. **(G)** The estimate of HSC frequency in BM for each experimental group is shown, as are the 95% confidence intervals for this estimate. **(H)** Pairwise analysis of experimental groups using Chi-squared test.

**Figure S4.**
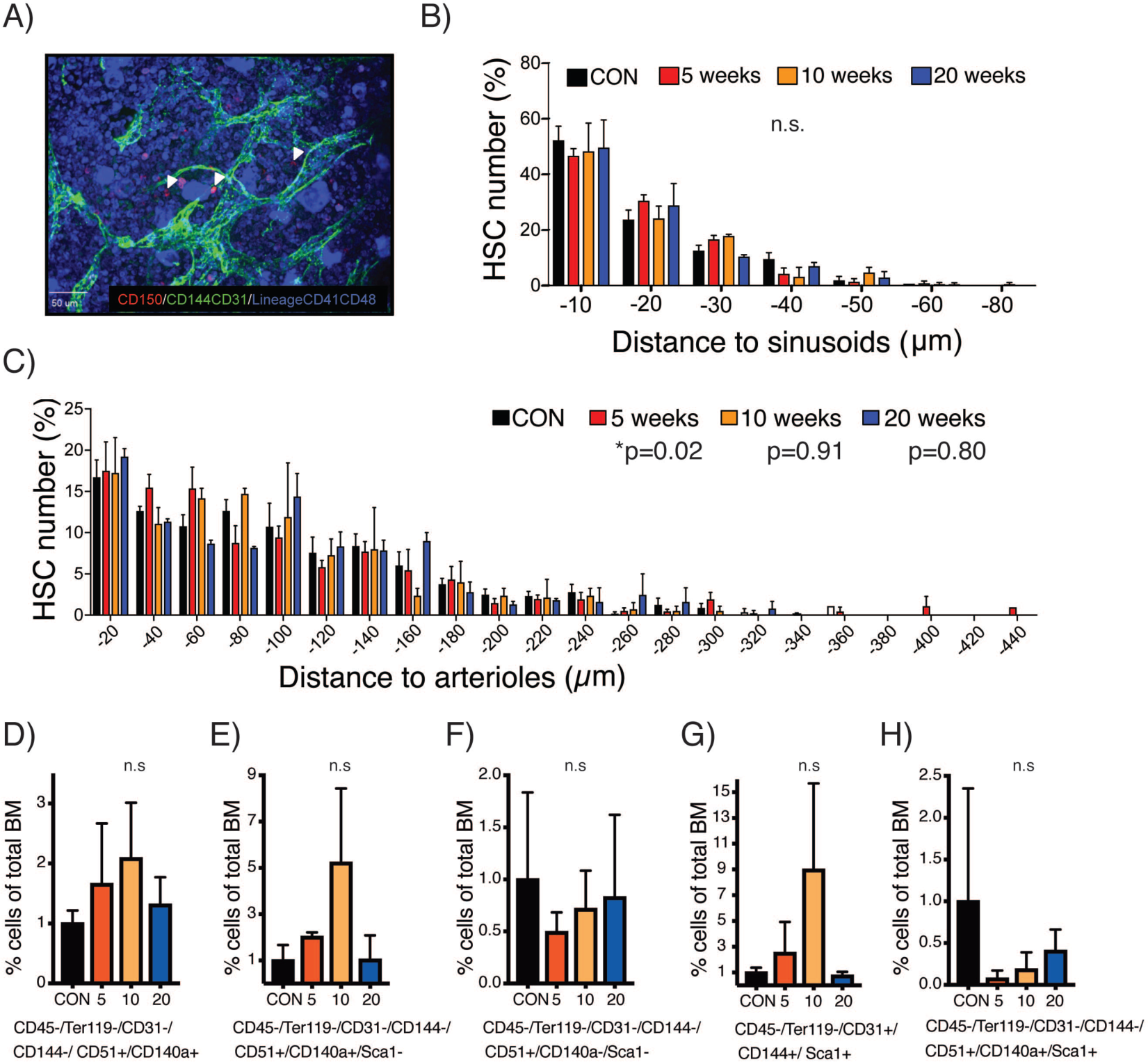
Repetitive exposure to inflammatory stress has a limited impact on BM niche architecture and composition. **(A)** Representative immuno-fluorescence image of a whole mount cross section of the sternum, used for the quantification of the spatial distribution of HSCs relative to the BM vasculature (sinusoids or arterioles). HSCs were identified by staining for CD150 expression (in red) and negativity for CD41 and CD48 lineage markers (in blue). HSCs are indicated by white arrowheads. Endothelial cells were defined by positivity for CD144 and CD31 expression (in green). **(B-C)** The spatial relationship between HSCs and the BM vasculature was quantified by measuring the shortest distance between HSCs and **(B)** sinusoids or **(C)** arterioles, n=4-5 mice, n=157-492 HSCs. Two-sample Kolmogorov-Smirnov tests were used for the comparison of distribution patterns between CON mice and the different recovery time points. **(D-H)** Analysis of the relative composition of defined niche components within the BM was performed using flow cytometry after staining with antibodies directed against a combination of CD45, Ter119, CD51, CD31, CD144, CD140α, Sca-1. Based on previous literature, these populations were analogous to **(D)** Nestin^+^ like stromal cells; **(E)** Cxcl12 abundant reticular; **(F)** osteoblast; **(G)** endothelial; and **(H)** mesenchymal stem cells. Mean ± SD are shown, n=3-9 mice. Statistical significance between control and treatment groups was evaluated by one-way ANOVA on ranks with Dunn’s multiple comparison tests (n.s=p>0.05).

**Figure S5.**
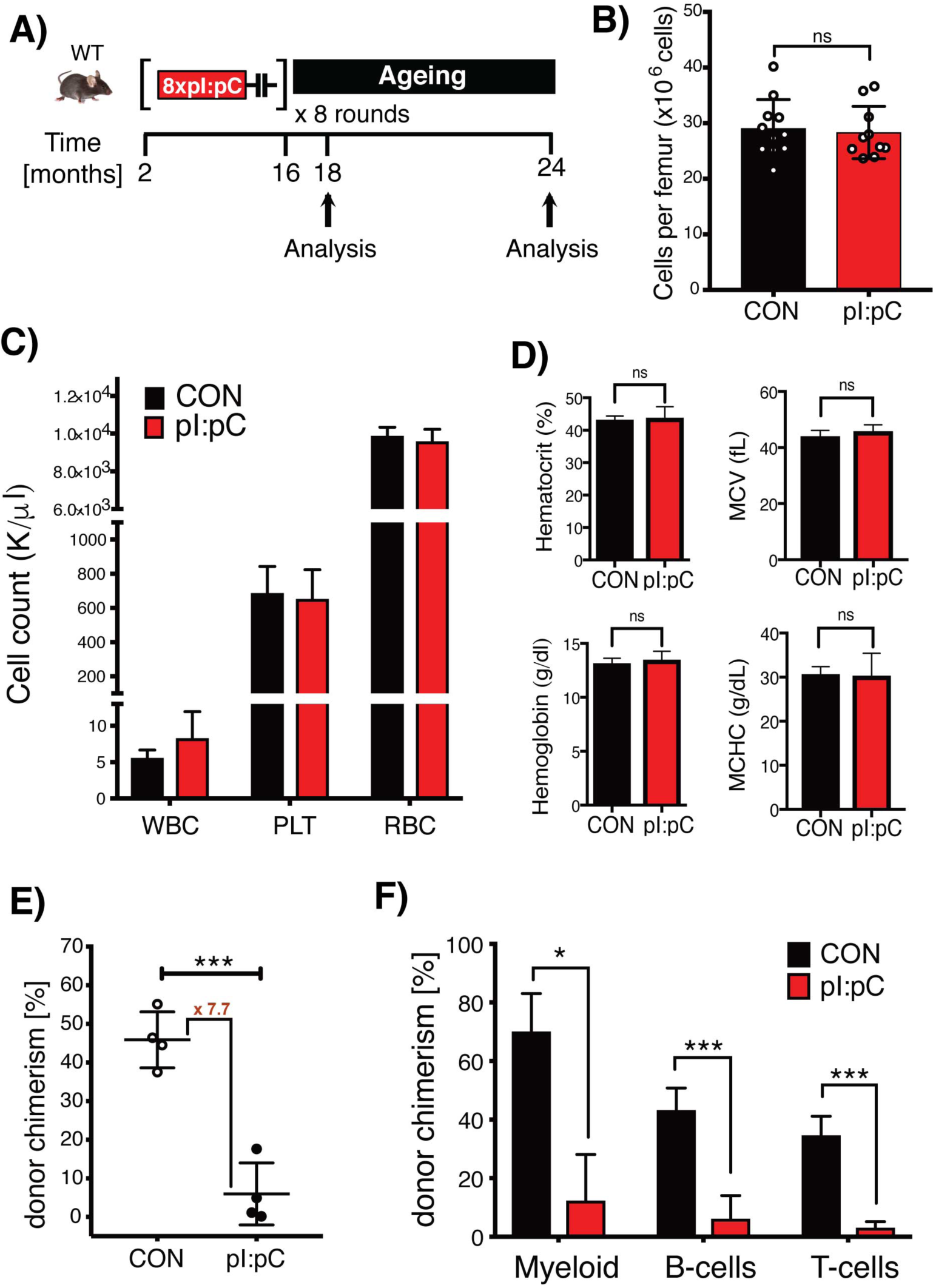
PB counts and BM cellularity at 18 months of age are not affected in mice exposed to repetitive to inflammatory stress, but HSCs demonstrate a profound functional engraftment defect at 24 months of age. **(A)** Schematic representation of the treatment schedule. C57BL/6J mice were treated for 8 consecutive rounds with pI:pC (64 injections). Controls (CON) represent either age matched non-treated mice, or age-matched mice that were subject to PBS injections in place of pI:pC. At the time of the last injection, the mice were 16 months old (68wk). They were then either sacrificed at 18 months of age and analyzed, or were left to age for an additional 8 months until they reached 24 months of age. **(B-D)** PB and BM parameters evaluated at 18 months of age. **(B)** BM cellularity **(C)** Numbers of leukocytes (WBC), platelets (PLT) and red blood cells (RBC) found in the PB. **(D)** Erythrocyte related parameters, including mean cell volume (MCV) and mean cell hemoglobin content (MCHC). Mean ± SD are shown, n=10-12 mice per group. **(E-F)** BM isolated from mice at 24 months of age was assessed for functional activity using a competitive repopulation assay, where test BM cells from individual donors were co-injected into recipients along with a 1:1 ratio of competitor BM cells. **(E)** Analysis of the contribution of test BM cells to donor PB leukocyte chimerism at 6 months post-transplantation demonstrates a 7.7-fold reduction in competitive repopulation activity in BM cells from pI:pC treated mice. **(F)** Analysis of the relative contribution of test BM to the myeloid (CD11b^+^Gr-1^+^), B-cell (B220^+^) and T-cell (CD4^+^/CD8^+^) lineages at 6 months post-transplantation. mean ± SD are shown, n=3-4 mice per group. Statistical significance was determined by unpaired two-tailed t-test (*P<0.05,***P<0.001).

**Table S1.**
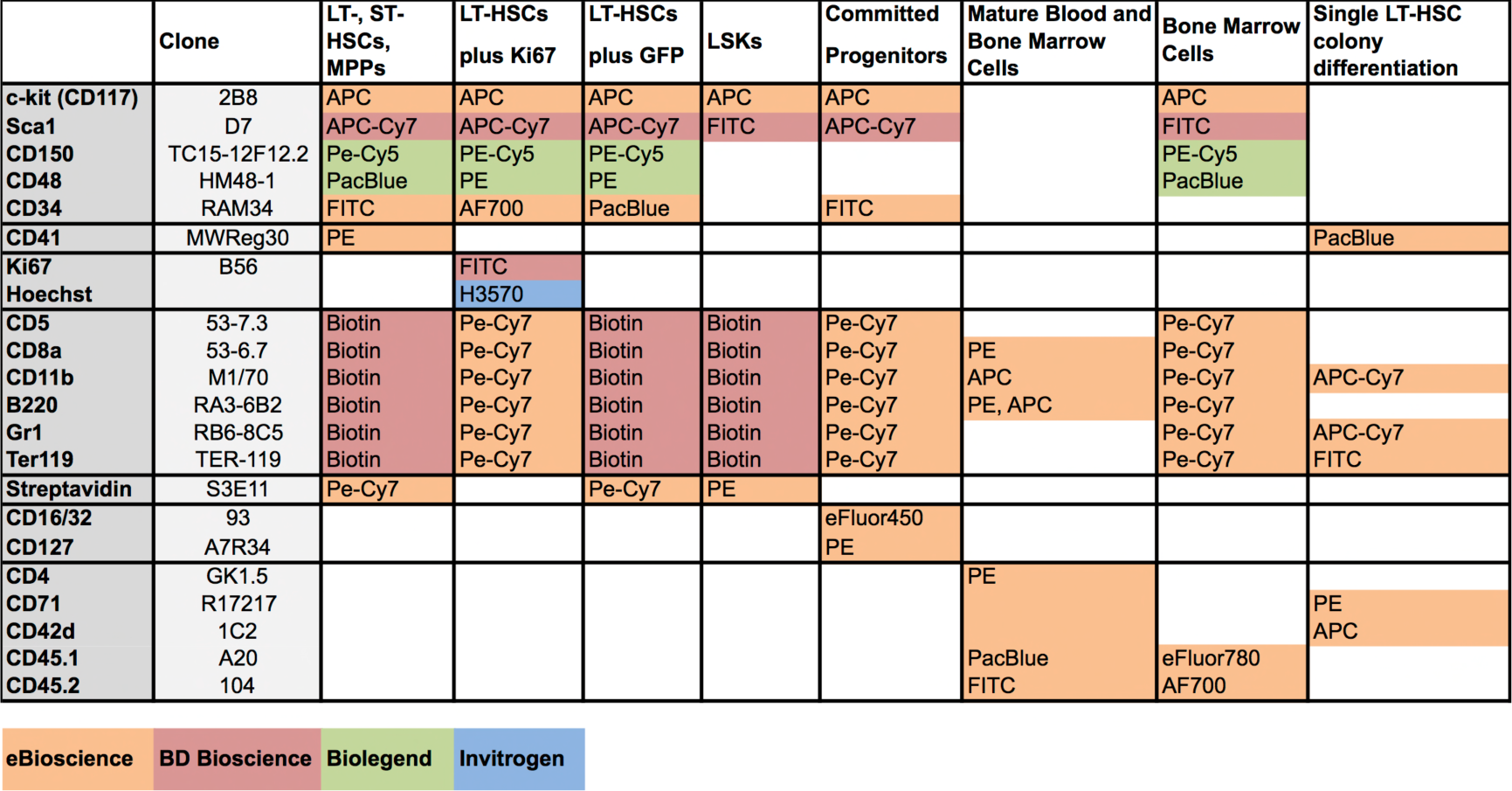
List of antibodies and antibody staining panels used in this study.

**Table S2.** Definition of specific immunophenotypically-defined cell populations e been analyzed in the manuscript.

| Abbreviation | Population | Surface markers |
| --- | --- | --- |
| LT-HSC | long term hematopoietic stem cell | Lin (-) c-kit(+) sca-1(+) CD150(+) CD48(-) CD34(-) |
| ST-HSC | short term hematopoietic stem cell | Lin (-) c-kit(+) sca-1(+) CD150(+) CD48(-) CD34(+) |
| HSC | hematopoietic stem cell | Lin (-) c-kit(+) sca-1(+) CD150(+) CD48(-) |
| MPP | multi potent progenitor | Lin (-) c-kit(+) sca-1(+) CD150(+) CD48(+) |
| CLP | common lymphoid progenitor | Lin(-)CD127(+);sca-1(mid) c-kit(mid) |
| MkP | megakaryocytic progenitor | Lin (-) c-kit(+) sca-1(-) CD150(+) CD41(+) |
| CMP | common myeloid progenitor | Lin (-) c-kit(+) sca-1(-) CD16/32(mid) CD34(+) |
| GMP | granulocytic-monocytic progenitor | Lin (-) c-kit(+) sca-1(-) CD16/32(hi) CD34(+) |
| MEP | megakaryocytic-erythroid progenitor | Lin (-) c-kit(+) sca-1(-) CD16/32(low) CD34(-) |
| KL | oligopotent progenitors | Lin (-) c-kit(+) sca-1(-) |
| Lin- | lineage negative cells | CD4(-) CD8(-) B220(-) Ter-119(-) CD11b(-) Gr-1(-) |
| Lin+ | lineage positive cells | CD4(+) CD8(+) B220(+) Ter-119(+) CD11b(+) Gr-1(+) |
| T-cell | mature T-cells | CD4(+) and/or CD8(+) |
| B-cell | mature B-cells | B220(+) |
| Myeloid | mature myeloid cells | CD11b(+) Gr-1(+) |
| CAR | Cxcl12 abundant reticular cell | CD45/Ter119(-) CD51(+) CD31(-) CD144(-) CD140alpha(+) sca-1(-) |
| OB | osteoblast | CD45/Ter119(-) CD144(-) CD51(+) CD31(-) CD140alpha(-) sca-1(-) |
| EC | endothelial cell | CD45/Ter119(-) CD31(+) CD144(+) sca-1(+) |
| MSC | mesenchymal stem cell | CD45/Ter119(-) CD51(+) CD31(-) CD140alpha(+) sca-1(+) |
| Nestin (+) like SC | Nestin(+) like stromal cells | CD45/Ter119(-) CD144(-) CD51(+) CD31(-) CD140alpha(+) |

